# Maternal Genetic Ancestry and Legacy of 10^th^ Century AD Hungarians

**DOI:** 10.1101/056655

**Authors:** Aranka Csősz, Anna Szécsényi-Nagy, Veronika Csákyová, Péter Langó, Viktória Bódis, Kitti Köhler, Gyöngyvér Tömöry, Melinda Nagy, Balázs Gusztáv Mende

**Author notes:** These authors contributed equally to this study (Aranka Csősz, Anna Szécsényi-Nagy). Present address: Department of Genetics, Hungarian Institute for Forensic Sciences, Budapest, Hungary (Gyöngyvér Tömöry).

## Abstract

The ancient Hungarians originated from the Ural region in today’s central Russia and migrated across the Eastern European steppe, according to historical sources. The Hungarians conquered the Carpathian Basin 895–907 AD, and admixed with the indigenous communities.

Here we present mitochondrial DNA results from three datasets: one from the Avar period (7^th^ –9^th^ centuries) of the Carpathian Basin (n = 31); one from the Hungarian conquest-period (n=76); and a completion of the published 10^th^-12^th^ century Hungarian-Slavic contact zone dataset by four samples. We compare these mitochondrial DNA hypervariable segment sequences and haplogroup results with published ancient and modern Eurasian data. Whereas the analyzed Avars represents a certain group of the Avar society that shows East and South European genetic characteristics, the Hungarian conquerors’ maternal gene pool is a mixture of West Eurasian and Central and North Eurasian elements. Comprehensively analyzing the results, both the linguistically recorded Finno-Ugric roots and historically documented Turkic and Central Asian influxes had possible genetic imprints in the conquerors’ genetic composition. Our data allows a complex series of historic and population genetic events before the formation of the medieval population of the Carpathian Basin, and the maternal genetic continuity between 10^th^- 12^th^ century and modern Hungarians.

## Introduction

According to historical sources, the Hungarian tribal alliance conquered the eastern parts of the Carpathian Basin in 895 AD, and in successive campaigns occupied its central territories until 907 AD ^1^. The mixed autochthonous population, which mostly spoke different Slavic, Turkic Avar, and German languages, integrated with variable speed with the newcomers, as we know from contemporaneous sources ^2^. Whereas the Slavs lived mainly on the fringes, the successors of the Avars persisted in some inner territories of the Carpathian Basin. The Avars arrived in the Carpathian Basin in 568 AD, fleeing the westward-expanding influence of the Turkic Khaganate in Inner Asia ^3^. The Avar population already included several folk elements at this time; and the population was uniform from neither a cultural nor a physical anthropological perspective. Over one hundred thousand excavated graves from the Avar period in the Carpathian Basin picture a heterogenic physical anthropological composition of this population, which contained mainly Europid characters and, only in certain regions and periods, was dominated by Asian craniometric indices ^4^. The occupation policy of Avar and ancient Hungarian tribes were similar due to similar steppe-type husbandry and management of space and power. In the politically unified alliance of the Hungarian tribes, both the leader and the tributary folks influenced each other culturally. These interactions are easily seen from the changing material culture of the Hungarian conquerors, who began to use local types of jewels but also maintained steppe-like traditions during the 10^th^ century ^5^. It is difficult to estimate the size of the 10^th^ –11^th^ century population of the Carpathian Basin from ca. twenty-five thousand excavated graves ^5,6^. Scholars estimate the Hungarian conqueror population in the Carpathian Basin between a few thousand and half a million, while the indigenous population size, which is also uncertain, is estimated at a few hundred thousand people ^7^.

Historical sources give evidence of the mixed ethnic composition of the Hungarians before the conquest of the Carpathian Basin ^2,8^. The diverse origin of the Hungarian tribes has also been documented in physical anthropological research. Craniometrical analyses revealed that the Europid crania type was predominant in the conquerors, with smaller amounts of Europo-Mongoloid characters ^9^. Regional groups of the ancient Hungarian anthropological series show morphometric parallels ranging from the Crimean Peninsula to the Kazakh steppe ^10^.

The Finno-Ugric origin of the Hungarian language is well recorded by linguistic research, which lead to an assumption that there was a Uralic substrate of the ancient Hungarian population ^2^. However, Turkic-speaking groups could also have had a significant role in the formation of the Hungarian people and political institutions, as suggested by ancient Turkic loanwords in the early layer of the Hungarian language and the Turkic origin of toponyms and person names of tribe leaders of the conquest-period ^11^. After leaving the Central Uralic homeland, an obvious source of the Turkic influence was the Turkic-speaking political environment of the Bulgars (Onogurs) and Khazars in the 9^th^- century Eastern European steppe, where the Hungarians lived for a period of time. The exact route and chronology of the Hungarian migration between the Ural region and the Carpathian Basin is continually debated among archaeologists, linguists and historians.

The genetic origin of ancient Hungarians is still in question, although some modern and ancient DNA studies have focused on this issue. For example, Tömöry et al. have described the mitochondrial DNA (mtDNA) of a small group of ancient Hungarians from the 10^th^ –12^th^ century Carpathian Basin, where the ancient Hungarians’ affinity to modern day Central Asia has been demonstrated. Tömöry et al. concluded, without simulation tests, that there was no genetic continuity between the classical conquerors and modern day Hungarians ^12^. A small 10^th^ –12^th^ century population from the northwestern Carpathian Basin has been reported with heterogeneous maternal genetic characteristics similar to modern Europeans ^13^. On the other hand, ancient mitochondrial DNA data from the putative source region of the ancient Hungarians is still scarce, and concentrates only on the prehistory of Siberia and Central Asia ^14–17^. Of four analyzed Y chromosomes from the conqueror population, two showed connections to Uralic peoples through N1c1 haplogroup marker Tat ^18^.

Genetic research of modern Hungarians has been a subject of four further mtDNA and Y chromosomal studies. Brandstätter et al. and Egyed et al. built the mtDNA control region and Y chromosomal STR databases from different groups of modern Hungarians, including an “average” Hungarian group from Budapest and two groups of Hungarian minorities – Ghimes Csango and Szekler – living in modern Romania. Both Szeklers and Csangos were found to harbor some Asian genetic components, and the Csango population shows genetic signs of long term isolation, which differentiated them from the Szeklers and the population of Budapest ^19–21^. Asian genetic mtDNA and Y chromosome components are apparently rare in the modern Hungarian gene pool, which led Semino et al. to the conclusion that the Hungarian conquerors were in small number and that the Hungarian language could be an example of cultural dominance ^22^. The pitfalls of the very hypothetical historical interpretation of modern day population genetic results have been critically reviewed by Bálint^23^.

The archaeogenetic contribution to the historical era of the Avar and conquest-periods (6^th^ –10^th^ centuries) in the Carpathian Basin is still sparse. Our research approaches the questions of maternal genetic composition and the origin of the ancient Hungarians, analyzing a dataset four times larger than previous work has attempted. The connections of the conquerors to the previous Avar and contemporaneous Slavic-Hungarian contact zone population will be determined, as well as connections to other ancient populations of Eurasia that have previously been published. We also compare our dataset with modern day data from the Carpathian Basin and Eurasia, in order to better understand the maternal genetic origin and legacy of the 7^th^ –11^th^ century population of the Carpathian Basin.

We focused on these questions through analysis of the mtDNA of 144 early medieval individuals from the 7^th^ –9^th^ centuries Avar and the 10^th^ –11^th^ century Hungarian period (Fig. 1).

**Figure 1.**
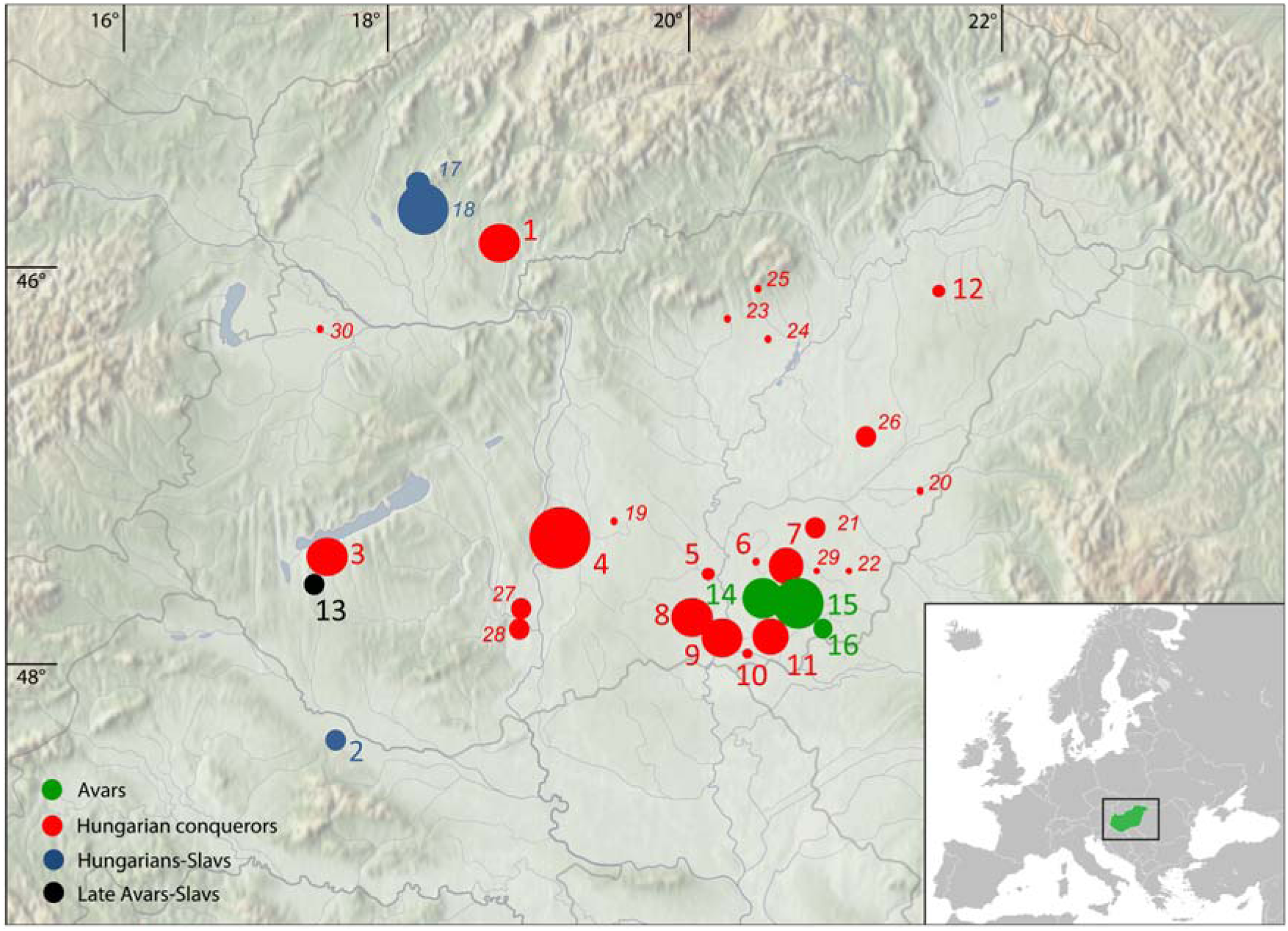
Location of investigated sites in the Carpathian Basin. Sizes of circles indicate number of obtained mtDNA haplotypes from a certain site. Italic letters (17–30) mark previously published data ^12,13^. Green color indicates Avar cemeteries, red color designates conqueror cemeteries, blue shows the contact zone, and black indicates 9^th^ –10^th^ century late Avar populations. Map was generated in Adobe Illustrator CS6 software. The map of Europe was downloaded from Wikipedia (https://commons.wikimedia.org/wiki/File%3ABlank_map_of_Europe.svg.) Numbers of successfully typed individuals are in brackets after site names: 1. Levice-Géňa (9); 2. Zvonimirovo (4); 3. Balatonújlak-Erdődűlő (10); 4. Harta-Freifelt (16+1); 5. Baks-Iskola (3); 6. Szentes-Borbásföld (1); 7. Szentes-Derekegyháza (8); 8. Kiskundorozsma-Hosszúhát (9); 9. Szeged-Öthalom (8); 10. Kiszombor (2); 11. Makó-Igási járandó (8); 12. Nyíregyháza-Oross Megapark (2); 13. Vörs-Papkert (5); 14. Szegvár-Oromdűlő (8 Avar+ 2 conqueror); 15. Székkutas-Kápolnadűlő (14); 16. Pitvaros-Víztározó (4); 17. Čakajovce (5); 18. NitraŠindolka (14); 19. Izsák-Balázspuszta (1); 20. Magyarhomorog (1); 21. Orosháza (1); 22. Szabadkígyós-Pálliget (1); 23. Aldebrő-Mocsáros (1); 24. Besenyőtelek-Szőrhát (1); 25. EgerSzépasszonyvölgy (1); 26. Sárrétudvari-Hízóföld (4); 27. Fadd-Jegeshegy (5); 28. MözsSzárazdomb (3); 29. Örménykút (3); 30. Lébény-Kaszás (1).

## Results

Reproduced hyper variable segment I (HVS-I) sequences were obtained from mtDNA of 111 individuals from the medieval Carpathian Basin: 31 mtDNA profiles from Avars, 76 from Hungarian conquerors and four from the southern Hungarian-Slavic contact zone (see Supplementary Table S3). The mtDNA of 111 individuals was extracted at least twice per individual from different skeletal elements (tooth and femur or other long bones, Supplementary Table S1), the HVS-I fragments were reproduced in subsequent PCR and sequencing reactions, at least twice per DNA extract. The sequence results of these replicates, spanning HVS-I nucleotide positions (np) 16040–16400, typing individual selections of 14 coding region positions and two fragments of the HVS-II (np 29-254) confirm the haplotypes to be authentic. Of the 144 processed samples, 33 had no amplifiable DNA yield, or the sequences gave ambiguous haplotype results.

The Avar group from the southeastern Great Hungarian Plain (Alföld) had a mixed European-Asian haplogroup composition with four Asian haplogroups (C, M6, D4c1, F1b) at 15.3%, but a predominantly European (H, K, T, U), haplogroup composition (Fig. 2). In the conqueror population the most common Eurasian haplogroups were detected. West-Eurasian haplogroups (H, HV, I, J, K, N1a, R, T, U, V, X, W) were present at a frequency of 77%, and Central and East-Eurasian haplogroups (A, B, C, D, F, G, M) at 23%. The most widespread haplogroups of the conqueror population were H and U with frequencies 22% and 20% respectively (Supplementary Table S5). Five individuals from the 9^**th**^ –10^th^ centuries from the west Hungarian Vörs-Papkert site were excluded from any statistical analysis because of their offside geographical location and cultural differences from the Avar and Hungarian sites. Their mtDNA belonged to the common European J and H haplogroups, but with rare haplotype variants in ancient and modern mtDNA databases (see Supplementary Table S15 for database references). The number of typed mtDNA from the 10^th^ –12^th^ century contact zone metapopulation ^13^ was enlarged by four 10^th^ century samples from present-day north Croatia. One belonged to a characteristic European H10e haplotype; another belonged to U7 haplotype, mainly distributed in modern Southwest Asia and Southern Europe; a third belonged to the Southwest Asian N1b1 type; the fourth U5a2a haplotype was common in modern Eurasia (private database, see Material and Methods, Supplementary Table S15).

**Figure 2.**
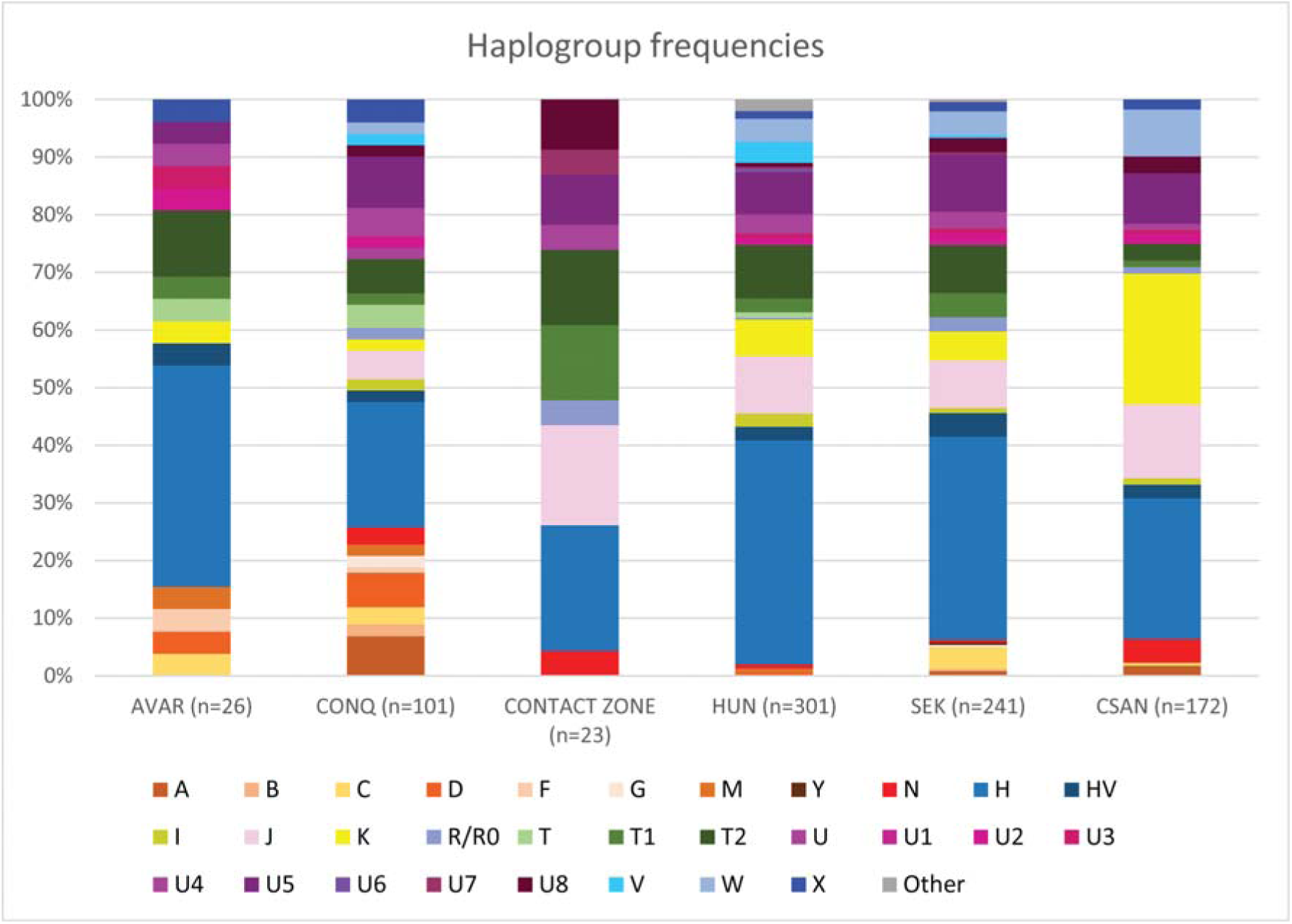
Haplogroup frequencies of the populations under study. Abbreviations: AVAR: Avars, CONQ: Hungarian conquerors, CONTACT ZONE: Hungarian-Slavic contact zone in the 10^th^-12^th^ centuries, HUN: modern Hungarians, SEK: modern Szeklers from Romania, CSAN: modern Csangos from Romania. The modern data were taken from ^12,20,49^ studies. For exact haplogroup frequencies see Supplementary table S5-S6.

The principal component analyses (PCAs) of ancient and modern-day populations were computed based on haplogroup frequencies (Supplementary Tables S5 and S6). PCA of 21 ancient populations showed a predominant difference between European and Asian populations, which indicates a clustering of the medieval populations of Europe, as well as the assembly of Avars, conquerors and further Mediterranean populations (Fig. 3A, Supplementary Fig. S1). Although the East Asian medieval populations were clearly separated from the European contemporaneous period on both PCA and Ward clustering, prehistoric Central Asian (Kazakhstan) and North Asian (Siberian Late Bronze Age Baraba) populations showed similarities to the conquest-period dataset in both analyses (Fig. 3B). The three Carpathian Basin populations were compared with populations from the most ancient North European and medieval Asian populations, showing significant differences in haplogroup composition (p<0.05). On the other hand, prehistoric Central Asian, south central Siberian (Minusinsk Hollow) and Baraba populations were not significantly different from the populations of the Carpathian Basin, and these affinities are also reflected in the clustering tree (Supplementary Table S5, Fig. 3B).

**Figure 3.**
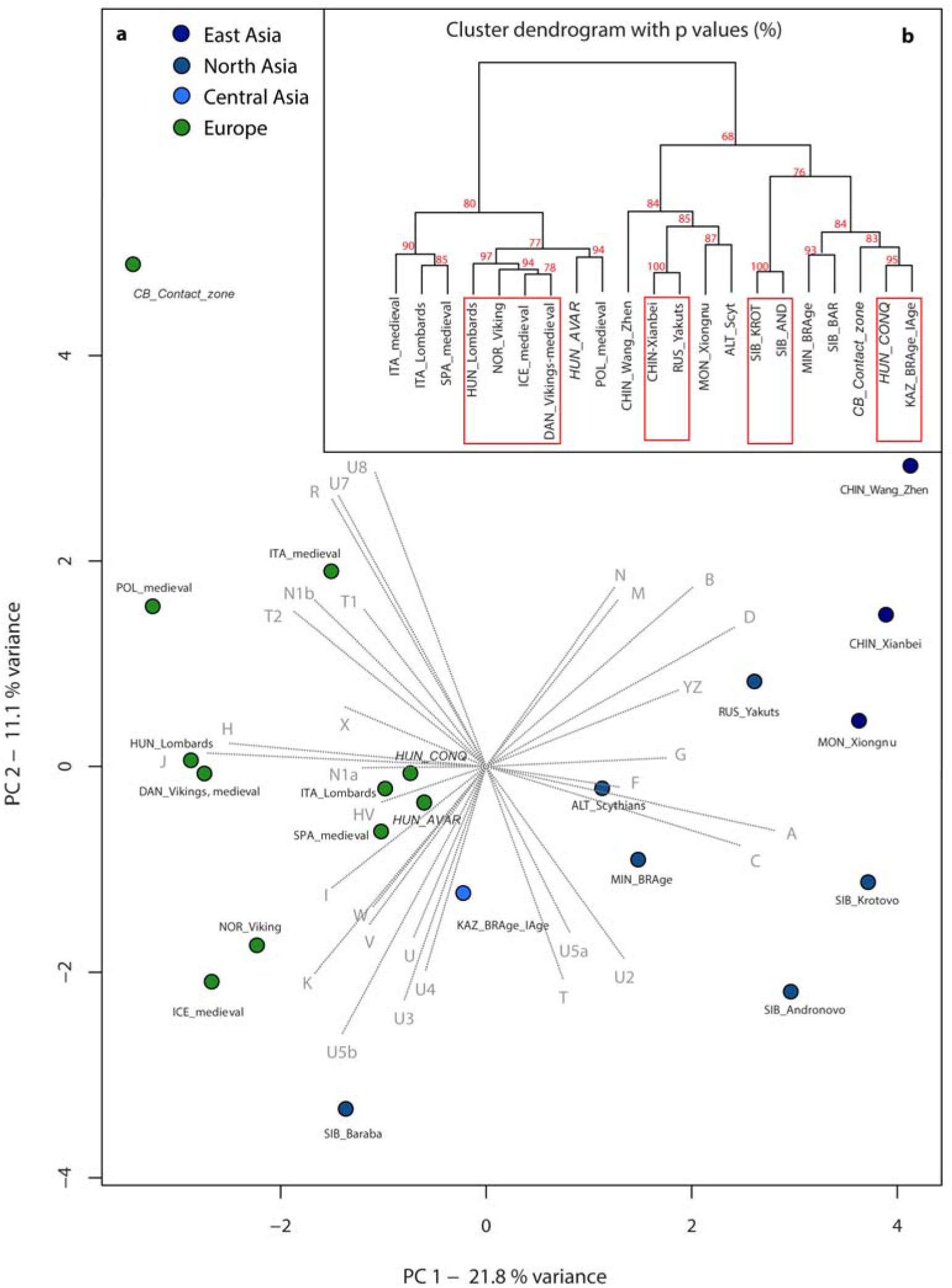
A: PCA plot of the first two components (32.9% of variance), comparing haplogroup frequencies of 21 ancient populations. B: Ward type hierarchical clustering of 21 ancient populations. On the PCA plot the contribution of each mtDNA haplogroup is superimposed as grey component loading vector. PCA of 21 ancient populations shows a predominant difference between European and Asian populations along PC1 (variance= 21.8%), which furthermore shows a clustering of the medieval populations of Europe, as well as the assembly of Avars (HUN_AVAR), conquerors (HUN_CONQ), and other Mediterranean populations. Along the PC2 component (variance = 11.1%), the most distant population within the European sector is of the contact zone in the Carpathian Basin (CB_Contact_zones). Prehistoric Central Asian (Kazakhstan), south western Siberian (Baraba Late Bronze Age culture), and south central Siberian populations (abbreviations: KAZ_BRAge_IAge; SIB_BAR, MIN_BRAge) show similarities to the conquest-period datasets both on PCA (A) and the Ward clustering tree (B). P values in percent are given as red numbers on the dendogram, where red rectangles indicate clusters with significant p-values. The abbreviations and references are presented in Supplementary Table S5.

The PCA of the investigated ancient and modern Eurasian populations demonstrated the clustering of most modern European populations by PC1, PC2, and PC3. Furthermore, their affinities to modern Near Eastern populations are represented by PC1 and PC3, whereas the modern Asian populations are dispersed along PC1. The conqueror population has a similar haplogroup composition to modern Central Asians and Finno-Ugric populations, which is also supported by Ward type clustering. While Avars rather showed modern European connections, the contact zone population had a Near Eastern type haplogroup composition (Supplementary Fig. S2, S3).

The distance calculations based on high subhaplogroup resolution also showed that modern Central Asian populations were highly similar to the conqueror population. The maternal genetic connections of the Avar group concentrated on modern Eastern European populations, and the contact zone group showed Southwest Asian affinities on genetic distance maps (GDM) (Fig. 6A, Supplementary Fig. S6A, S7A, see Supplementary Table S13 for references).

The haplogroup frequency-based test of population continuity (TPC) ^24^ rejected neither the null hypothesis of population continuity between the Avars and the southeastern Alföld group of conquest-period Hungarians, nor between Avars and all conquerors analyzed from the Carpathian Basin. Furthermore, the haplogroup frequency differences between the 10^th^-12^th^ century populations and modern Hungarians, and also Hungarian minorities of Szeklers and Csangos living in Romania can be explained by genetic drift that occurred during the last millennium (Supplementary Table S7).

Pairwise genetic distances were calculated between 21 ancient and 52 modern populations. Interestingly, pairwise F_ST_ values of Avars indicated non-significant differences among nearly all medieval European populations, and from Central Asia, as well as from many modern-day Europeans. The Hungarian conqueror population showed the lowest distances from modern-day Uzbeks and Turkmens (F_ST_= 0.00335 and 0.00489 respectively) and from six ancient populations: medieval Poles (F_ST_ = −0.00018), Bronze and Iron Age in present-day Kazakhstan (F_ST_ = −0.00164), Bronze Age along the south central Siberian flow of Yenisey River (Minusinsk Hollow) (F_ST_ = −0.00208), Siberian Baraba population (F_ST_ = −0.01003), Avars (F_ST_ = 0.00233), and 6^th^ century Lombards from Hungary (F_ST_ value 0.00762), these values are non-significant (p > 0.05). The distances from the ancient populations are visualized on an F_ST_ level plot (Fig. 4). The mixed contact zone population has the shortest distances from present-day Iraq (F_ST_ = 0.00817), Italy (F_ST_ = 0.00923), Czechs (F_ST_ = 0.01023) and Avars (F_ST_ = 0.01094). For the genetic F_ST_ values and their corresponding p-values, see Supplementary Table S8–S9.

**Figure 4.**
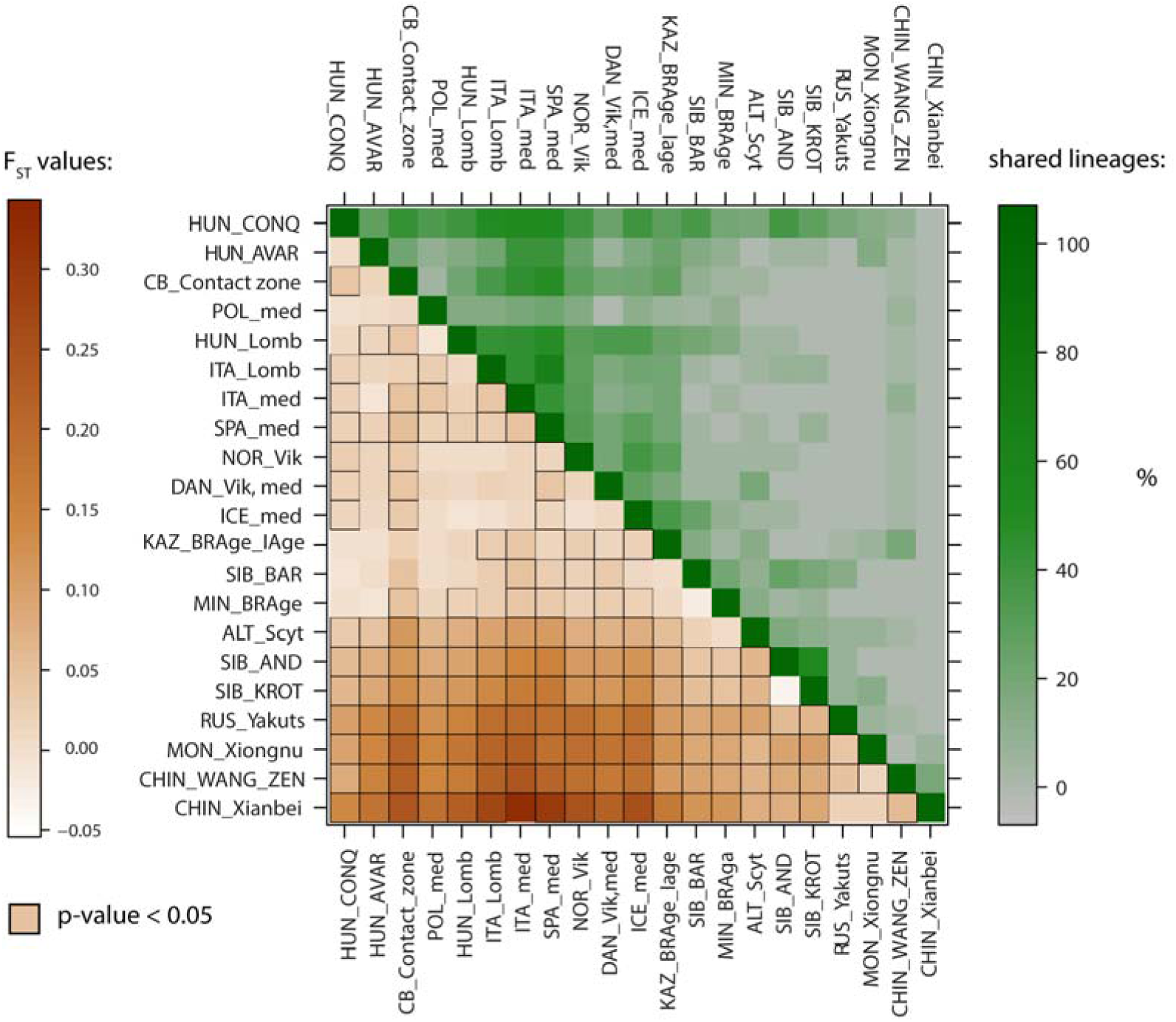
F_ST_ level plot and shared haplotype analysis with 21 ancient populations. Lower left corner: larger pairwise F_ST_ values indicating greater genetic distances are marked by dark brown shades. Significant p-values are highlighted with black squares. Upper right corner: high percentages of shared lineages are highlighted with dark shades of green color. For exact values, abbreviations and references, see Supplementary Table S8, S10.

In order to visualize these genetic distances, linearized Slatkin F_ST_ values were displayed on a multi-dimensional scaling (MDS) plot (Fig. 5, Supplementary Fig. S5 and Table S8–S9). The plot of ancient populations reflects the PCA and shows the connection between the south western Siberian Baraba population ^17^, south central Siberian Minusinsk Depression and Kazakhstani prehistoric populations ^14,16^ and the conquerors. The Avar and contact zone populations show stronger affinities to the European medieval populations, similarly to the PCA results. On the modern population MDS plot, which also contains the three investigated medieval datasets, a very similar picture is observable to the modern PCA, except that the Southwest Asian populations do not separate from Europe along coordinate 2 (Supplementary Fig. S5).

**Figure 5.**
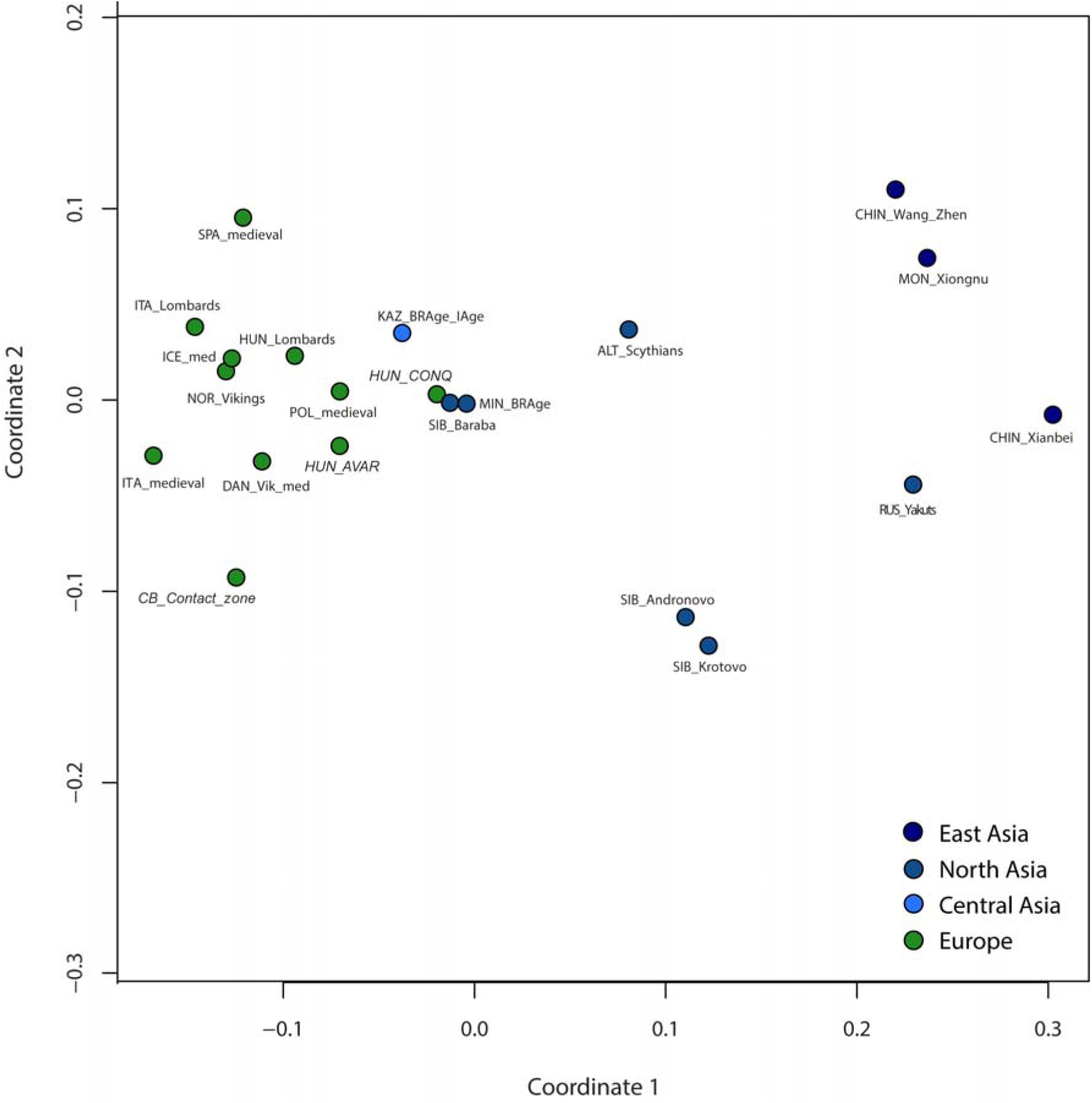
MDS with 21 ancient populations. Stress value is 0.08633. In the Slatkin F_ST_ calculation a 329 bp long fragment of the HVS-I was considered. The MDS plot of ancient populations shows the connection of the Siberian Baraba population (SIB_Baraba), Kazakhstan’s, and the south central Siberian Minusinsk Depression’s Bronze Age populations (KAZ_BRAge_IAge; MIN_BRAge) to the conquerors (HUN_CONQ). The Avars (HUN_AVAR) and contact zone population (CB_Contact_zone) show stronger affinities to the European medieval populations than the conquerors (Supplementary Table S8).

The sequence-based genetic distance maps, encompassing 141 modern populations, show congruently the Central Asian affinity to the conquerors, the European/Near Eastern characteristic populations to the Avar sequences, and predominant Near Eastern affinities to the contact zone group (Fig. 6B, Supplementary Fig. S6B, S7B, see Supplementary Table S14 for references).

**Figure 6.**
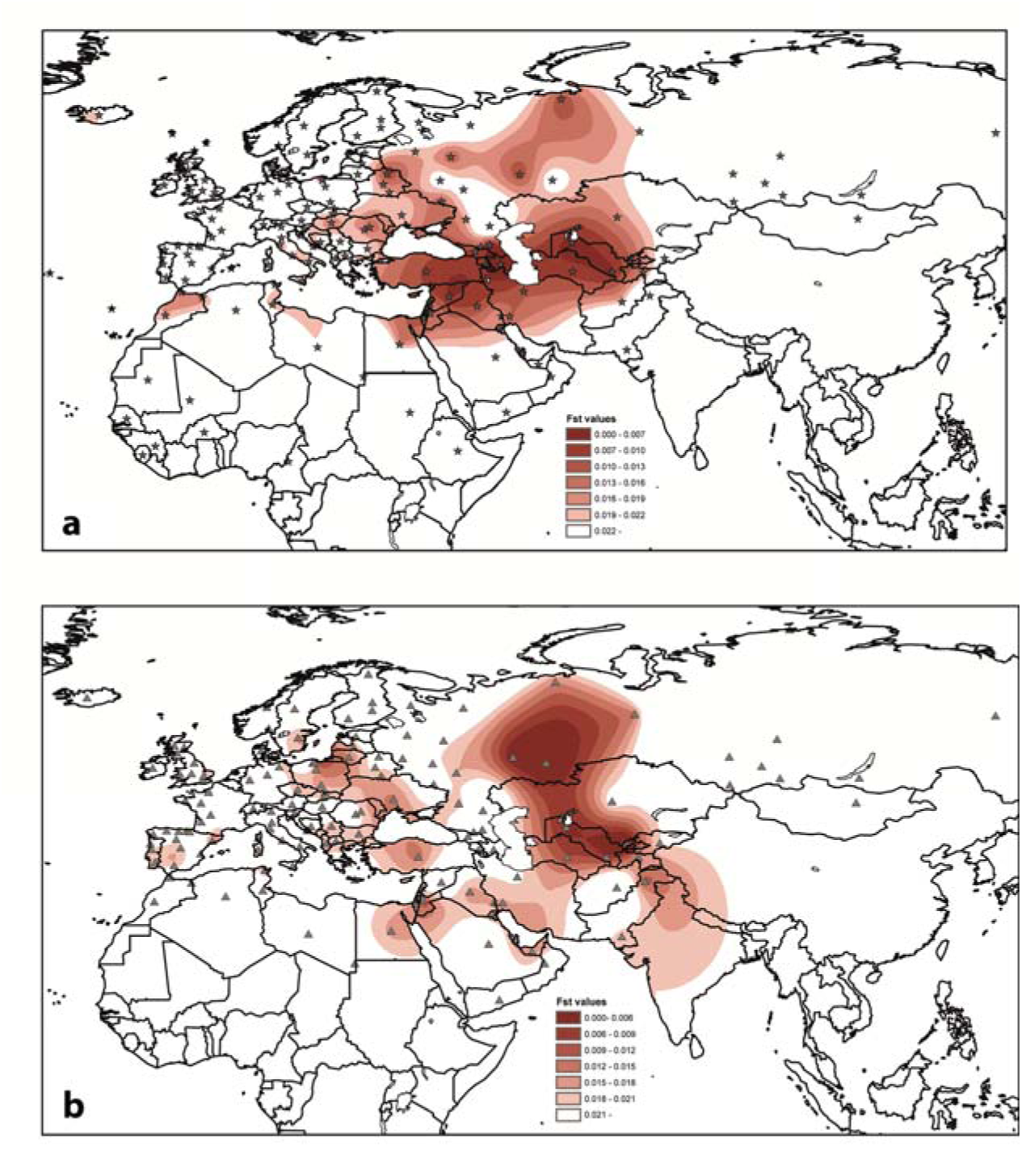
Mapping of genetic distances counted from the Hungarian conqueror population. A: Haplogroup frequency based genetic distances, B: HVS-I sequence based distances. Grey stars (A) and triangles (B) signalize the sampled and compared populations. White regions display F_ST_ values greater than 0.022 (A) or 0.021 (B) or unconsidered territories. (A) The genetic distance map based on high resolution haplogroup frequency of 157 modern populations shows that modern Central Asian populations are highly similar to the conquerors. It presents low distances between present-day Azerbaijan, North Caucasian District, Uzbekistan and some Near-Eastern populations. The values are presented in Supplementary Table S13. (B) The sequence based genetic distance map, encompassing 141 modern populations, shows the Central Asian affinity to the conquerors (with the highest similarity toward today’s Uzbekistan, the Russian population of the Bashkortostan Republic and the Tatar population of Russian Tatarstan). The values of genetic distances are listed in Supplementary Table S14. The F_ST_ values and coordinates were interpolated with the Kriging method implemented in Arcmap ArcGIS version 10.3 (https://www.arcgis.com).

The 101 ancient Hungarian samples belong to 75 HVS-I haplotypes (haplotype diversity Hd = 0.987). The haplotype diversity is highest in the Avar group, and lowest in the contact zone dataset (Table 1). The shared haplotype analysis (SHA) shows that medieval populations from Southern Europe (Spain and Italy) shared over 50% of haplotypes with the conqueror population (Fig. 4, and Supplementary Table S10). High proportions of shared lineages with the conquerors were detected in the contact zone population (43.5%), Vikings from Norway (39.3%), Iceland (39.7%), and 6^th^-century Lombards in Hungary (39.3%). The SHA analysis is strongly influenced by altering haplotype diversity and the high number of Cambridge Reference Sequence (rCRS) H lineages in medieval Spanish, Italian and Norwegian Viking groups, which causes high proportion of lineage sharing with only a small number (n = 4–5) of shared lineage types. Medieval populations from Italy and Spain shared many of their haplotypes (40–48%) with the Avar and contact zone populations as well. On the other hand, many lineages of the Bronze Age Andronovo, Baraba, and Bronze Age population of the region of today’s Kazakhstan were shared with the conquerors (37.5–29.4%), with some identical Asian lineages among them.

**Table 1.** Diversity indices. Abbreviations: n means number of HVS-I sequences, h means number of haplotypes; Hd means haplotype diversity; Pi means nucleotide diversity; k means average number of nucleotide differences. Diversity indices were tested for np 16040-16400.

|  | n | h | Hd | Pi | k | Tajima's D | p values |
| --- | --- | --- | --- | --- | --- | --- | --- |
| Conqueror Hungarains | 101 | 75 | 0,987 | 0,01663 | 5,71928 | -2,01256 | <0.05 |
| Avars | 26 | 24 | 0,994 | 0,01456 | 5,25538 | -1,88457 | <0.05 |
| Hungarain-Slavic contact zone | 23 | 19 | 0,98 | 0,01676 | 5,73123 | -1,65039 | >0.05 |

We analyzed more deeply the sharing of the Eastern Eurasian haplotypes – found in the Carpathian Basin medieval datasets – with modern and ancient populations (Supplementary Table S11). Based on our updated Eurasian mtDNA database of 64,650 HVS-I sequences, the Asian lineages in the conqueror dataset showed diverse hits. Three Asian A haplotypes had no matches in our modern-day mtDNA database (see references in Supplementary Table S15). Other A11 and A12a haplotypes have parallels in present-day Uzbekistan, Kazakhstan, other Asian populations, in people of the Xiongnu confederation of the 3^rd^ BC to 2^nd^ AD century, in the late medieval Yakuts, and in medieval Scandinavia. Two B haplotypes were present in today’s China, Kazakhstan and spread as far as Thailand. The detected conqueror C-C4, F1b, and G2a haplotypes were widespread in modern Eurasia, and had parallels even in China and Korea. Six of these C, F, and G2a haplotypes had parallels in ancient populations of Asia. Among the five Asian D lineages, two were unique in the database and two were common in Central and East Asia. One D haplotype however (Der4.522) showed rare occurrence in Kazakhs, Uzbeks and Altaians, and Siberian populations. Among the Avars, three Asian haplotypes (C, M, D4c1) were found. One C haplotype had only one match in modern Kazakh population, the other M lineage was common in Central and East Asia, but also occurred in Southwest Asia and Europe. The third Asian haplotype is D4c1, which also occurred at low frequency in Central, North, and East Asia (Supplementary Table S11). It is noted that other lineages belonging to Western Eurasian type haplogroups could also be brought into the Carpathian Basin from Central/North Asia, for example, U4 or T types that were also frequent in ancient and modern Siberia ^17^.

We selected 23 modern populations from the GDM, MDS, and PCA datasets, which possibly had increased lineage-sharing with the conquerors and we compared them using a modern SHA (Supplementary Table S12). Populations speaking Uralic languages are not well studied for mtDNA, therefore we could only use Khantys, Mansis, Nenets, and Komis as references for Uralic peoples. The ancient conquest-period population had the highest lineage-sharing with the Tatars in Russian Tatarstan, and the Nenets and Komi groups (42–36%). They are followed by Hungarians, Russians in Bashkortostan, and three populations of almost identical percentages; Ukrainians, the Khanty and Mansi population, and Szeklers. When counting lineages, rather than the number of sequences, Csangos, Khantys and Mansis, and the population of the Russian Bashkortostan Republic were the third, fourth and fifth populations with the highest lineage-sharing (22.6–17%). Interestingly, the relatively low lineage sharing with Uzbeks and Turkmens did not reflect the high similarities visible on MDS and GDMs (Fig. 6, Supplementary Fig. S5).

## Discussion

We typed the mtDNA of 111 medieval individuals and performed population genetic and statistical analyses, focusing on three populations that existed in the 7^th^ –12^th^ centuries in the Carpathian Basin. The earliest population under study is the 7^th^ –8^th^ century Avars from the southern part of modern Hungary (Fig. 1). The genetic results from the Avars demonstrate their predominant southern and eastern European maternal genetic composition, with some Asian elements. The local continuity of the Avar population on the southern Great Hungarian Plain to the Hungarian conquest-period cannot be rejected by haplogroup based simulation analyses (TPC, Supplementary Table S7) and was also demonstrated on PCA plots (Fig. 3a, Supplementary Fig. S1). However, sequence-based tests and shared haplotype analyses showed a low level of identical maternal lineage among the Avars and ancient Hungarians, even when including the geographically connecting southeastern group of the conquerors in the calculations (Fig. 4, Supplementary Table S10). The Avar dataset originates from a single micro regional group of the complex Avar society, who buried their dead in catacomb graves (26). Furthermore, anthropological results showed that this part of the Avar population represents mostly Europid, local morphological characters, and therefore it cannot be used as a proxy of the whole Avar population of the Carpathian Basin. Further regional groups should be analyzed from the late Avar period for a better estimation of the Avar-Hungarian continuity.

The Hungarian conqueror genetic dataset from the 10^th^ century showed more explicit connections toward Central Asian ancient and modern populations, in contrast to the preceding Avars. Asian haplogroups occurred among both male and female conquerors (Supplementary Table S1, S3), which can be an argument for a Hungarian settlement in which both men and women took part. It reflects the physical anthropological and archaeological data, which showed that, not only an armed population stratum, but a whole population arrived in the Carpathian Basin ^25^. However, Asian lineages in the conqueror dataset can also be an argument for the continuity of the Avars, who could have mixed and acculturated during the Hungarian conquest-period ^26^. We would need more Avar period genetic data, especially from the late Avar period to assess this hypothesis.

In a previous study, Tömöry et al. presented mitochondrial genetic data of 26 Hungarian conquerors, who were divided into “commoners” (n = 15) and “high status” (n = 12) groups according to the excavated grave goods ^12^. The latter group shows more heterogeneous haplogroup composition, and also some haplotypes that are rare in modern populations. We do not follow this concept in our current study, because grave goods cannot represent evidence of social status with a high level of certainty ^26,27^, and therefore levels of richness or status cannot be categorized precisely. Furthermore, people of low social status could also have been part of the conqueror community, who most probably arrived from the east of the Carpathian Basin as well. Chronological subdivision of the studied graves is also challenging, even ^14^C dating is not accurate enough for dating 9^th^ –10^th^ centuries AD.

Most of the Asian mtDNA lineages occurred in 10^th^ century cemeteries with small numbers of graves (7–18 graves), and identical lineages were found among cemeteries, rather than within them. This is especially interesting in light of the fact that seven analyzed cemeteries have been completely excavated (Kiskundorozsma, Balatonújlak, Harta, Makó-Igási járandó, Levice-Géňa, Szeged-Öthalom, Szentes-Derekegyháza graveyards). This phenomenon suggests that the conquerors had a mobile way of life or can be explained by the strong marriage connections of the Hungarian communities. The lack of, or small number of intra-cemetery maternal relations is striking at the sites Kiskundorozsma and Levice-Géňa (nine typed and maternally unrelated individuals in both cases), Szeged-Öthalom (eight unrelated people) and Harta. At the Harta site, fifteen women, three men and two children were excavated. We found only one pair of females with identical HVS-I sequences (a common rCRS H type), but in other cases the maternal kinship relation among the 16 typed individuals could be excluded by HVS-I analyses. Many academic archaeologists explain that the small conqueror graveyards are small family graveyards, and use the grave goods of the assumed generations in these graveyards as chronological horizons ^28^. The example of Harta raises the possibility that family relations were not the sole rule of burial order. Mobile groups of people could use these cemeteries for a short period of time. These observations are relevant for the relative chronological and socio-archaeological assumptions about the 10^th^ century Carpathian Basin. Nevertheless, other classic 10^th^- century graveyards, such as Balatonújlak, contained more signs of possible maternal relations within the cemetery (Supplementary Table S3). The unequal geographic distribution of the samples did not allow us to make further conclusions on the internal (geography or chronology based) genetic structure of the presented 10^th^- century population of the Carpathian Basin.

We found genetic similarities of the conquerors with the Late Bronze Age population of the Baraba region, situated between the rivers Ob and Irtis ^17^, and with Bronze Age and Iron Age populations that lived in central Asia ^15^ and south Siberia ^14,16^. Comparing the conqueror mtDNA dataset to a large modern-day population dataset, we also found comprehensive genetic affinities towards modern populations of Central Asia and Central Russia. The parallels of these Asian haplogroups are found in modern ethnic groups speaking both Ugric and Turkic languages. The historically and linguistically assumed homeland of the ancient Hungarians was in the Central Ural region, which is an easily accessible part of the mountain range. Finno-Ugric-speaking groups might have settled on both sides of the Urals during the early Medieval period ^29^. Archeological records, for example, from central-eastern Uralic site Uelgi, indicate archaeological cultural mixture of northern Ugric and eastern steppic Turkic elements. These eastern components show cultural connections toward the region of the Emba River in today’s western Kazakhstan and toward the Srostki culture ^30^, which indicates that the ancient Hungarian population could already have been reached in the Central Ural region by several cultural and genetic influences. Newly revised archaeological connections of the Central Urals and the Carpathian Basin suggest a quick migration from the forest steppe to the Carpathian Basin ^31^, and during these events, the genetic make-up of the conquerors retained some Central Asian signatures.

Modern-day Hungarians were very similar to their surrounding Central European populations from the maternal genetic point of view, as is demonstrated by previous mtDNA studies ^12,19^. In our analyses, the Hungarian speaking Szekler, Ghimes, and Csango minorities in today’s Romania showed differing genetic connections from each other. Whereas the Szekler population was consistent with the Central and Eastern European maternal genetic diversity, the haplogroup and haplotype composition of the Csangos was more related to Near Eastern populations (Supplementary Fig. S4, Table S9). These results correspond to the fact that the Csangos, in the Romanian Ghimes region, are a genetically isolated population ^20^, living separately from both Romanians and Szeklers.

The maternal gene pool of Csangos, Szeklers and “average” Hungarians can be descended from 9^th^ –11^th^ century ancient Hungarians, and the differences in their haplogroup composition from the conquerors can be explained by genetic drift (Supplementary Table S7). It is an interesting phenomenon that some Asian haplogroups (A, B, C, G2a) that occur in the conquerors also occurred among Szeklers. This could suggest a sizeable legacy of the conquerors or it may mean that these Asian influences reached Romania in other time periods. Of the 76 detected conqueror haplotypes, 21 had matches in the modern Szekler and Hungarian populations (11.2–15.4% of all lineage types), but none were Asian (Supplementary Table S11). Fourteen conqueror lineages had matches in the Csango dataset, which represents a greater proportion (22.6%) of the total number of Csango lineage types, one of which belonged to the Asian C haplogroup. We would need more medieval samples from Romania and a reconsidered sampling of the current population in the Carpathian Basin in order to better estimate the genetic relations among past and present populations.

The 10^th^ century population of the Carpathian Basin had regionally different, but mostly heterogeneous physical anthropological and linguistic natures, which could be a consequence of the varied ethnic and linguistic composition of the conquerors. On the one hand, this parallels with the genetic diversity of the conquerors, and that the tribe alliance of the Hungarians was a culturally and linguistically mixed community in the steppe ^2^. On the other hand, it could also be a consequence of the mixture of several populations, which had experienced the conquest-period in the Carpathian Basin and the geopolitical environment of the new homeland. The mixed nature of the newly founded Hungarian State was documented in the early 11^th^ century, and described as a basic characteristic of a successful medieval state ^11^. The samples from the 10^th^ –12^th^ century contact zone dataset from the fringes of the Hungarian territory originate from different geographic regions. They represent a mixed dataset within medieval Europe, which showed haplogroup-level connections to the conquerors and ancient Asia (Fig. 3B), but on the sequence level they had affinities with medieval Poles, Lombards, and Avars. Their subsisted maternal genetic signature was found today in Southern Europe and the Near East (Fig. S2, S7). Written sources document the diverse acculturation speed of local populations in the Carpathian Basin. For example, the population of the Čakajovce settlement slowly adopted items of Hungarian traditions to their culture ^32^. This process could last 100–150 years, until burials with poor costume elements and jewels appeared, and Christian cemeteries became used. A new mixed culture began to form in the mid-10^th^ century, which disseminated in the whole territory of the Hungarian Principality regardless of ethnicity.

The results presented here provide a picture of the maternal gene pool of three medieval populations in the Carpathian Basin. Research should continue with the analysis of whole mitochondrial genomes for more exact haplogroup definitions, and Y chromosomal genetic diversity of these populations, in order to define the paternal genetic components of these populations, along with possible sex differences in migration and dispersal patterns. Furthermore, genome-wide sequencing of these samples and analyses of the comparatively ancient (early medieval) Eastern European, Central and North Asian data, which are currently still lacking, might reveal further signs of origin and admixture of the populations discussed here. Moreover, this may shed light on a complex population genetic structure of the first millennium BC of West, North, and Central Eurasia.

## Conclusion

This study contributes ancient mtDNA data to the research on Hungarian ethnogenesis and the conquest-period. We present the first described Avar-period ancient DNA dataset (n = 31), an almost four-fold enlargement of the existing Hungarian conquest-period dataset (with n = 76), and a magnified dataset from the Hungarian-Slavic contact zone of the 10–12^th^ centuries (with n = 4). These together with the previously published 10^th^-12^th^ century results were compared with published ancient and modern Eurasian mtDNA data. The results comprehensively demonstrate the conqueror maternal gene pool as a mixture of West Eurasian and Central/North Eurasian elements. Both the linguistically recorded Finno-Ugric roots and the Turkic, Central Asian influxes had possible genetic imprints in the conquerors’ mixed genetic composition. The small number of potential intra-site maternal relations compared to the number of detected inter-sites relations suggests that conqueror communities were mobile within the Carpathian Basin. Our data support the complex series of population genetic events before and during the formation of the 10^th^- century population of the Carpathian Basin. These processes might be defined by future ancient DNA studies focusing in the Ural region and in the Eastern European steppe using genome-wide sequencing techniques.

## Materials and Methods

### Sample information, ancient DNA work

The human skeletal remains (bones and teeth) used in this study were collected from 6^th^ –10^th^ century cemeteries excavated in the Carpathian Basin. The sampling was performed by co-workers of the Institute of Archaeology, considering various aspects: (1) geographical location; (2) chronology; (3) archaeological characteristics; (4) grave goods ^33^.

We investigated 144 medieval samples: from the Hungarian conquest-period 88 samples were analyzed from the cemeteries of Harta-Freifelt, Balatonújlak-Erdődűlő, Kiskundorozsma-Hosszúhát, Baks-Iskola, Szeged-Öthalom, Makó-Igási járandó, Szentes-Derekegyháza, Nyíregyháza, Kiszombor, Szentes-Borbásföld (all from Hungary), and Levice-Géňa (Slovakia). Furthermore, four Avar-period cemeteries were studied with 50 samples collected from Szegvár-Oromdűlő, Pitvaros-Víztározó, Székkutas-Kápolnadűlő, the 9^th^ –10^th^ century Vörs-Papkert (all from Hungary), and six samples from one site in the medieval Hungarian-Slavic contact zone, Zvonimirovo (located in present-day northern Croatia). Nine samples from these sites were already part of G. Tömöry’s PhD dissertation ^34^ (Fig. 1, Supplementary Table S1). It is important to note that the graves of the conquest-period population are mainly dated to the 10^th^ century. They were probably not the first generation of conquerors, which is very problematic to distinguish at the current state of research. One further point to note is that the Avar samples belong to a single micro region of the Avar Khaganate, and therefore do not represent the whole Avar population of the Carpathian Basin.

Sampling was carried out using gloves, facemasks, and body suits, in order to minimize the risk of contamination by contributors. Two bone fragments, usually two compact bone tissues from different parts of long bones, or one tooth and one compact bone fragment of a femur were collected from each individual. All stages of work were performed under clean conditions in a dedicated ancient DNA laboratory at the Institute of Archaeology, Research Centre for the Humanities, Hungarian Academy of Sciences, Budapest, following published ancient DNA workflow protocols and authentication criteria ^12,13,35^. Laboratory rooms for pre-PCR and post-PCR works were strictly separated. All pre-PCR steps (bone cutting, surface removing, powdering, extraction, PCR set-up) were carried out in separate clean rooms. The laboratory work was carried out wearing clean overalls, facemasks and face-shields, gloves and over-shoes. All materials and work areas were bleached and irradiated with UV-C light. We used PCR-clean plastic wares and Milli-Q ultrapure water for reaction preparation. In order to detect possible contamination by exogenous DNA, one milling blank per sample, one extraction and amplification blanks per every five samples were used as negative controls. MtDNA haplotypes of all contributors (anthropologists, geneticists) in the sampling and laboratory work were determined in the post-PCR lab, and compared with the results obtained from the ancient bone samples. Only one haplotype match was found between an ancient sample (PitV124.436B) and an anthropologist, who had no contact with this specific sample (Supplementary Table S16).

The specimens were prepared following the protocols described by Kalmár et al. ^36^ and Szécsényi-Nagy et al. ^37^. The bone and teeth samples were bleached, washed, and irradiated with UV-C light (1.0 J/cm^2^, 25 min). The surfaces of teeth samples were cleaned by sandblasting (Bego, EasyBlast), while the surfaces of bone samples were removed with a fresh drilling bit at slow speed, followed by UV exposure for 30 min on each side. Bone and tooth pieces were mechanically ground into fine powder in a sterile mixer mill (Retsch MM301).

Different DNA extraction methods were used, repeatedly validating the results per sample ^12,36,38^. MtDNA hypervariable segment I (HVS-I) and coding region positions were amplified in several PCRs in a total volume of 40 μl reaction mix, containing 5 μl DNA extract, 1×AmpliTaq Gold-Buffer; 2 U AmpliTaq Gold DNA polymerase (Applied Biosystems); 200 µM of each of the dNTP; 25 pmolμl^−1^ primer; 1.5 mM MgCl_2_; 4mgml^−1^ BSA. The HVS-I region of mtDNA was amplified in two overlapping fragments with two sets of primers, and an additional 16 primer pairs were used to amplify haplogroup-diagnostic nucleotide positions in coding regions (see Supplementary Table S2). Cycling parameters were 98°C for 10 min; followed by 39 cycles of denaturation at 98°C for 30 s, annealing at 56°C for 1 min, and extension at 72°C for 40 s; and a final step of 72°C for 5 min. PCR products were checked on 8% native polyacrylamide gel. The PCR products were purified using QIAquick^®^ PCR Purification Kit (Qiagen) following the manufacturer’s protocol, or purified from 2% agarose gel with Bioline Isolate PCR & Gel Kit in a final volume of 15 μl. Sequencing reactions were performed using the ABI PRISM BigDye Terminator v3.1 Cycle Sequencing Ready Reaction Kit (Applied Biosystems) and sequencing products were purified by ethanol precipitation. The sequences were determined on ABI PRISM 3100 (PE Applied Biosystems) in cooperation with BIOMI Ltd (Gödöllő, Hungary). The sequences were evaluated with Chromas Lite 2.4.1 and GeneDoc software ^39^.

The sequence polymorphisms in the nucleotide position range 16040–16400 were compared with the revised rCRS ^40^ as well as the Reconstructed Sapiens Reference Sequence (RSRS, www.mtdnacommunity.org) ^41^. Sequences were submitted to GenBank under the accession numbers KU739156–KU739266. Haplogroup determination was carried out according to the mtDNA phylogeny of PhyloTree build 17, accessed 18 February 2016 ^42^, and these haplogroup definitions were checked in our mtDNA database of 78,000 samples (enlarged database of that reported in ^24^), and in EMPOP.

We could not determine the haplogroup classification of one sample (HAR1.56B), due to detection failure of U haplogroup-diagnostic at coding region position 12308. Therefore we included it only into shared haplotype analyses (SHA) of HVS-I sequences, and excluded it from other statistical analyses.

### Reference population data

Of the typed 111 mtDNA profiles, we excluded the site Vörs-Papkert from the population genetic analyses, because it represents a 9^th^ –10^th^- century late Avar-Slavic mixed population in Transdanubia (western-Hungary). On the other hand, we included 26 samples from medieval Hungary described by Tömöry et al.^12^, and 19 samples from medieval Slovakia ^13^ into the population genetic analyses because of their similar historical, chronological and geographical traits to the new sample sets. We created three groups from a total of 150 analyzed Carpathian Basin samples for population genetics analyses: (1) conquest-period dataset (75 new samples and 26 samples described by Tömöry et al., 2007); (2) Avars in the southeastern part of today’s Hungary (26 samples) (3) “contact zone” (23 samples of the conquest-period derived from the outskirts of medieval Hungary: 4 new samples from today’s Croatia and 19 samples from the cemeteries of Nitra-Šindolka and Čakajovce (today’s Slovakia) described by Csákyova et al. ^13^ (Supplementary Table S4).

The ancient datasets were compared with 57,098 published modern HVS-I sequences as well as 614 medieval sequences of European, Near Eastern and Asian populations: Lombards from Hungary and Italy, medieval population from northern Italy, medieval Basques from Spain, medieval populations from Poland, Iceland and Denmark, Vikings from Norway and Denmark, three ancient populations (3^rd^ century BC–14^th^ century AD) from Mongolia and Inner Mongolia (China), and late medieval Yakuts from Russia. In addition, in order to have a proxy for the genetically uncharacterized first millennium AD populations of Central and North Asia, we used prehistoric (Bronze Age and Iron Age) datasets from modern-day Russia, Kazakhstan and Mongolia. Their characteristics, abbreviations and references are described in Supplementary Tables S5, S6 and S15.

### Population genetic analyses

Standard statistical methods were used for comparisons and calculations of genetic distances between our investigated populations (conquerors, Avars, and contact zone) and a further 18 ancient and 53–157 modern populations. Diversity indices were calculated in DNASP v5 ^43^ using sequence range np 16040–16400.

PCAs were carried out based on mtDNA haplogroup frequencies. We considered 31 mtDNA haplogroups in PCA of 21 ancient populations, while in PCA with the 3 medieval populations and 53 modern-day populations, 36 mtDNA haplogroups were considered (Supplementary Tables S5 and S6). All PCAs were performed using the prcomp function for categorical PCA, implemented in R 3.1.3 (R Foundation of Statistical Computing, 2015) and plotted in a two-dimensional space, displaying the first two or the first and third principal components, respectively (Fig. 3A, Supplementary Fig. S1–S3).

Hierarchical clustering was performed using Ward type algorithm ^45^ and Euclidean measurement method, where frequencies of the PCA haplogroups were used. The result was visualized as a dendrogram with the pvclust library in R.2.13.1 ^44^ (Fig. 3B). Cluster significance was evaluated by 10,000 bootstrap replicates. Significance of each cluster was given as an AU (Approximately Unbiased) p-value, in percentage. Fisher tests based on absolute haplogroup frequencies used in ancient PCA (except that U1 and Y, Z remained separated) were performed using sqldf library and fisher.test function in R.3.1.3.

Population comparisons were estimated using Arlequin 3.5.1 ^46^. Pairwise F_ST_ values were calculated based on 35,203 modern and 764 ancient HVS-I sequences (nucleotide positions (np) 16050–16383) of 83 populations: 21 ancient and 52 modern-day populations from Eurasia. Tamura & Nei substitution model ^47^ was assumed with a gamma value of 0.325 and 10,000 permutations were used for p-value calculation (Supplementary Table S8–9). The F_ST_ values were analyzed using MDS and applied on the matrix of linearized Slatkin F_ST_ values ^48^ (Supplementary Table S8–9) and visualized in a two-dimensional space (Fig. 5) using the metaMDS function based on Euclidean distances implemented in the vegan library of R 3.1.3 ^44^.

We tested the continuity of populations as described by Brandt et al. ^24^ with an absolute frequency of 22-37 mtDNA haplogroups. We performed tests assuming three effective population sizes (Ne = 500; 5,000; 500,000), and compared Avars with all conquest-period Hungarians, and with the southeast group of the latter (n = 45), who lived on the territory of the preceding Avar group. We also compared 10^th^-12^th^ century and modern-day Hungarians and the culturally isolated minority populations, Szekler and Csango, who live in Romania (Supplementary Table S7).

The shared haplotype analysis was carried out in order to detect and compare the mtDNA haplotypes shared between 21 Eurasian ancient populations, and to observe lineage sharing between the conquerors and 23 modern Eurasian populations. Identical HVS-I sequences and numbers of different lineage types were counted (Supplementary Table S10, S12). Asian lineages in the conqueror and Avar datasets were also counted in our database of 64,650 Eurasian sequences (Supplementary Table S11).

The comparative modern mtDNA datasets with detailed information on geographic origin were used for the GDM. From these datasets, we performed genetic distance calculations in two ways. First, we used high resolution haplogroup frequency tables of 157 populations (n = 49,439 individuals), differentiating 211 sub-haplogroups. We calculated genetic distances of these modern populations from the three Carpathian Basin medieval populations (Supplementary Table S13). Second, we randomly chose maximum 140 sequences per population (n = 18,499 sequences altogether), in order to balance the differences in sample sizes, and calculated F_ST_ values between medieval Carpathian Basin and 141 present-day populations. The sequence length was uniform, ranging np 16068–16365 (Supplementary Table S14). The analysis was performed in Arlequin software, using Tamura & Nei substitution model ^47^, with a gamma value of 0.177. For the haplotype definition, the original definition was used. F_ST_ values between conquerors, Avars, and the contact zone population and each modern population were combined with longitudes and latitudes according to population information in the literature. The F_ST_ values and coordinates were interpolated with the Kriging method implemented in Arcmap ArcGIS version 10.3.

## Acknowledgement

We are grateful to Csanád Bálint and István Raskó for the establishment of the Laboratory of Archaeogenetics in the IA RCH HAS, and their help in initializing the genetic research on Hungarian ethnogenesis. We thank Gábor Lőrinczy, József Szentpéteri, Csilla Balogh, Attila Türk, Attila Jakab, Gábor Nevizánsky for providing the human samples. We also thank Dóra Kis’ contribution to the laboratory work, and Antónia Marcsik and Erika Molnár for providing samples and anthropological data. The authors are also grateful to Guido Brandt for collecting the modern reference data.

## Author contributions

BGM, PL, MN designed the study. AC, VB, GT performed ancient DNA analyses. VC, ASN performed the population genetic analyses. KK and BGM carried out the anthropological analyses. ASN, VC, PL, BGM wrote the paper. All authors read and commented the manuscript.

